# Porcine cytomegalovirus detection by nanopore-based metagenomic sequencing in a Hungarian pig farm

**DOI:** 10.1101/2022.12.28.522123

**Authors:** Adrienn Gréta Tóth, Regina Fiam, Ágnes Becsei, Sándor Spisák, István Csabai, László Makrai, Tamás Reibling, Norbert Solymosi

## Abstract

The rapid diagnosis of infectious diseases has an essential impact on their control, treatment and recovery. Oxford Nanopore Technologies (ONT) sequencing opens up a new dimension in applying clinical metagenomics. In a large-scale pig farm in Hungary, four fattening and one piglet nasal swab pooled samples were sequenced using ONT for metagenomic analysis. Long reads covering 53.69% of the porcine cytomegalovirus genome were obtained in the piglet sample. The 650 bp long read matching the *glycoprotein B* gene of the virus is sequentially most similar to Japanese, Chinese and Spanish isolates.

## Introduction

In both human and animal health, the rapid diagnosis of diseases, including infectious diseases, has a crucial impact on their treatment and recovery. Oxford Nanopore Technologies (ONT) sequencing opens up a new dimension in the application of clinical metagenomics^1^ in veterinary medicine^2^. This third-generation sequencing technique can rapidly provide information on specific samples’ microbial components in a few hours to some days. While the preceding and parallel NGS (next-generation sequencing) methodologies provide higher sequence detection reliability, the sequencing time does not allow rapid microbial diagnostics in practice.

The results presented here are from a study aimed at gaining experience in the clinical metagenomic applicability of ONT in veterinary medicine. Here, we present only the main virological result of veterinary relevance: the detection of porcine cytomegalovirus (PCMV) sequences. The infection occurs in almost all pig populations, but clinical disease is rare, except in young piglets, where it can be fatal.^3^ However, since xenotransplantation from PCMV-infected pigs affects the recovery and survival of human patients, screening these donor animals has become an important issue.^4–6^

Although the virus has already been identified by PCR in Hungary^7^, a phylogenetic comparison of its sequence has not yet been performed. In addition to ONT-based viral detection, the similarity of its genome sequence to the available genomes is studied in this work.

## Materials and Methods

Nasal swab samples were collected from 5-week-old piglets of the same stable and from 16 (sample id: 3, 4) and 19-week-old (id: 1, 2) fattening pigs of two-two boxes of two stables from a Hungarian large-scale swine farm located near the town of Szekszárd on 21 November 2022. After sample collection, the nasal swabs were transported on ice and stored at -20 °C before the laboratory procedures. Porcine nasal swabs were pooled in nuclease-free molecular biology water as follows. Each five fattening pig samples deriving from the same stable and box, and two piglet pools of four-four piglet samples were created.

### DNA extraction and metagenomics library preparation

DNA extraction was performed with QIAamp Fast DNA Stool Mini Kit from Qiagen. The concentrations of the extracted DNA solutions were evaluated with an Invitrogen Qubit 4 Fluorometer using the Qubit dsDNA HS (High Sensitivity) Assay Kit. The concentrations of the 2 piglet samples were insufficient for library preparation. Thus, the two extracted piglet-deriving DNA solutions were pooled and concentrated with a vacuum concentrator. Consequently, the library preparation was conducted on DNA deriving from four fattening pig nasal swab samples and one piglet nasal swab sample. The metagenomic long-read library was prepared by the Ligation Sequencing Kit (SQK-LSK110) combined with the PCR-free Native Barcoding Expansion 1-12 (EXP-NBD104) from ONT. The sequencing was implemented with a MinION Mk1C sequencer using an R9.4.1. flow cell from ONT.

### Bioinformatic analysis

From the generated FAST5 files, one fast (configuration file: `dna_r9.4.1_450bps_fast_mk1c.cfg`) and one high-accuracy (configuration file: `dna_r9.4.1_450bps_hac_mk1c.cfg`) base calling was performed by ONT’s Guppy basecaller (v6.4.2, https://nanoporetech.com/community). The further analytical steps were done using the two-way called sequences parallel. The raw reads were adapter trimmed and quality-based filtered by Porechop (v0.2.4, https://github.com/rrwick/Porechop) and Nanofilt (v2.6.0)^8^, respectively. The resulting reads were taxonomically classified using Kraken2^9^ with the NCBI non-redundant nucleotide database^10^. Evaluating the taxon hits, the cleaned reads were mapped to the reference genome of *Suid betaherpesvirus 2* (KF017583.1) by minimap2.^11,12^ The sequence matching the *glycoprotein B* gene (*gB*) was used in the phylogenetic analysis. For that purpose, we used the available (19/12/2022) sequences with complete or partial *gB* CDSs. By blastn (BLAST v2.13.0+)^13^ with default settings, pairwise alignments were performed to identify the matching range and strand of the subject sequences. By the MUSCLE aligner (v3.8.1551)^14^, multiple sequence alignment was performed on the cropped subject and the query sequences. To construct a maximum-likelihood tree^15^, the function pml, optim.pml (`model=‘JC’`, `optNNi=TRUE`, `optBf=TRUE`, `optQ=TRUE`, `optInv=TRUE`, `optGamma=TRUE`, `optEdge=TRUE`) from the phangorn (v2.10) package was applied.^16^ All data processing and visualization were performed in the R environment.^17^

## Results

In total, 5.94 and 5.97 gigabases of data were generated during the 72 hours of sequencing, according to fast and high-accuracy basecalling. The descriptive statistics of the sequences generated by the two basecalling procedures are summarized in Figure 1. The high-accuracy basecalling generated slightly more nucleotides and longer reads, while the number of reads was equal.

**Figure 1.**
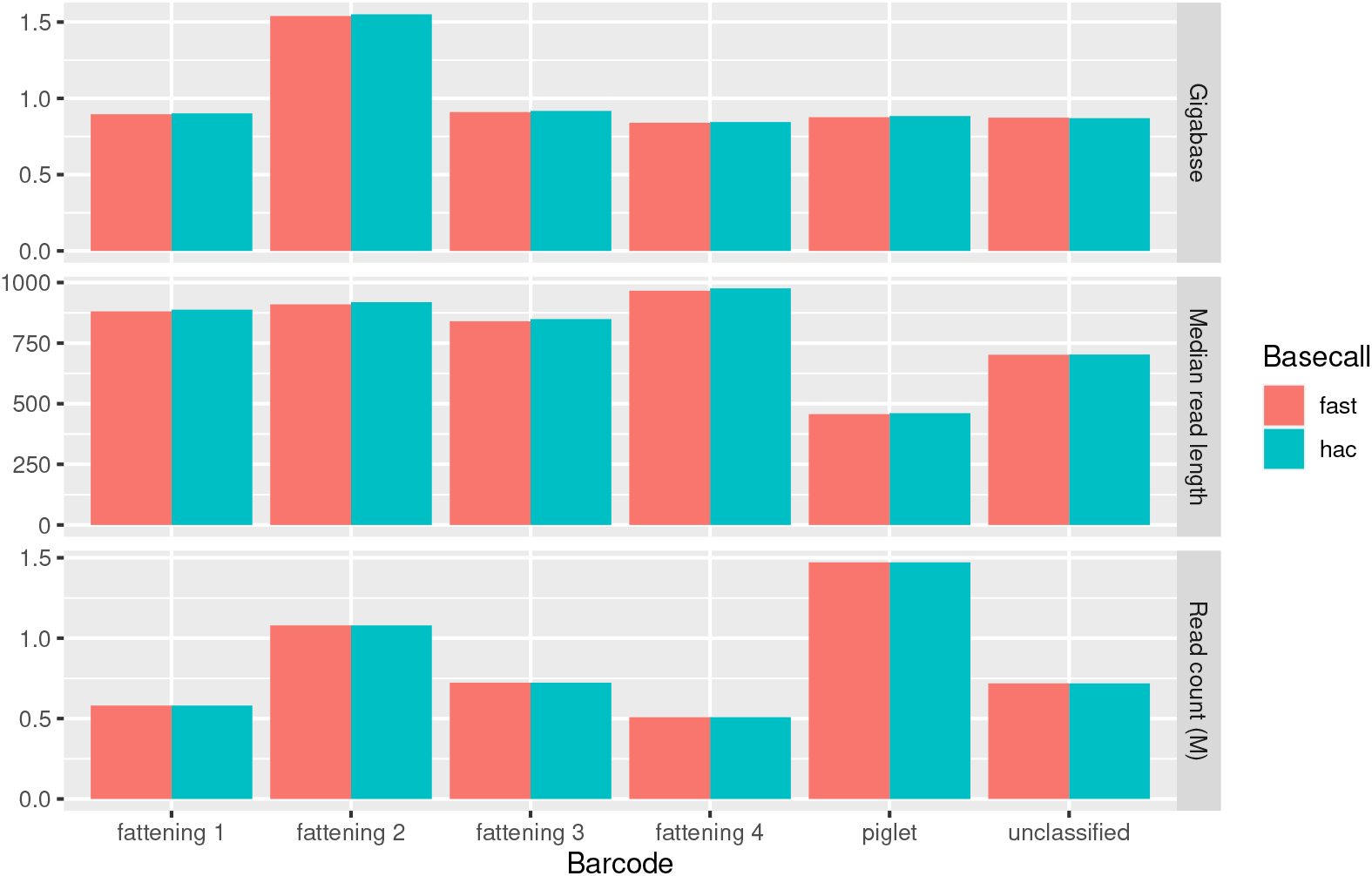
Descriptive statistics of raw reads. The high-accuracy (hac) basecalling, compared to the fast one, generated slightly more nucleotides and longer reads, while the number of reads was equal.

In the sample fattening 1, 2, 3, 4, the matched read number on the *Suid betaherpesvirus 2* reference genome was 2, 5, 8, 1, respectively. None of the four fattening samples had a read that matched *gB*. Of the unclassified (without barcode) reads, 25 matched the genome, and no hit was on gene *gB*. In the piglet sample, 315 reads aligned to the reference genome, covering 53.69% of it.

One sequence overlaps the *gB* gene and covers the region between nucleotides 49507 and 50172 of the representative reference genome. Where the identity was 631/671 (94%) and 641/665 (96%) with gaps 32/671 (4%) and 18/665 (2%) for the fast and high-accuracy basecalled sequence, respectively. The high-accuracy called sequence similarity with other *gB* sequences is shown by cladogram in Fig 2. Based on sequence similarity, the multiple sequence alignment of the closest strains is shown in Fig 3. The deletions identified in our sample are not found in the other *gB* genes. In addition to the deletions, only two positions (insertions at 87 and 139) of the sequential variations are shown in Fig 3, where no other *gB* gene modifications were found.

**Figure 2.**
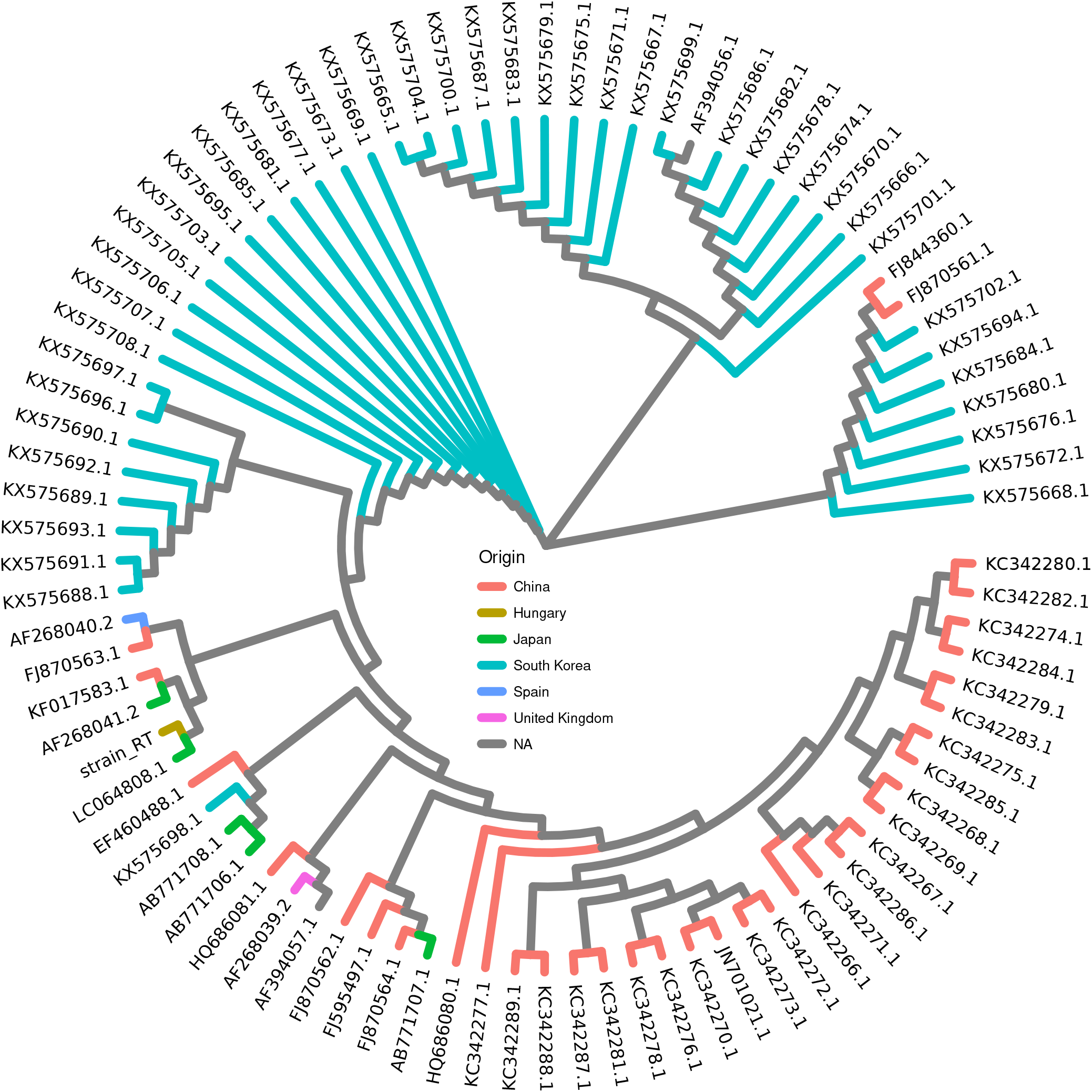
Cladogram based on *gB* gene sequence similarity. The `strain_RT` is the sequence obtained by high-accuracy basecalling.

**Figure 3.**
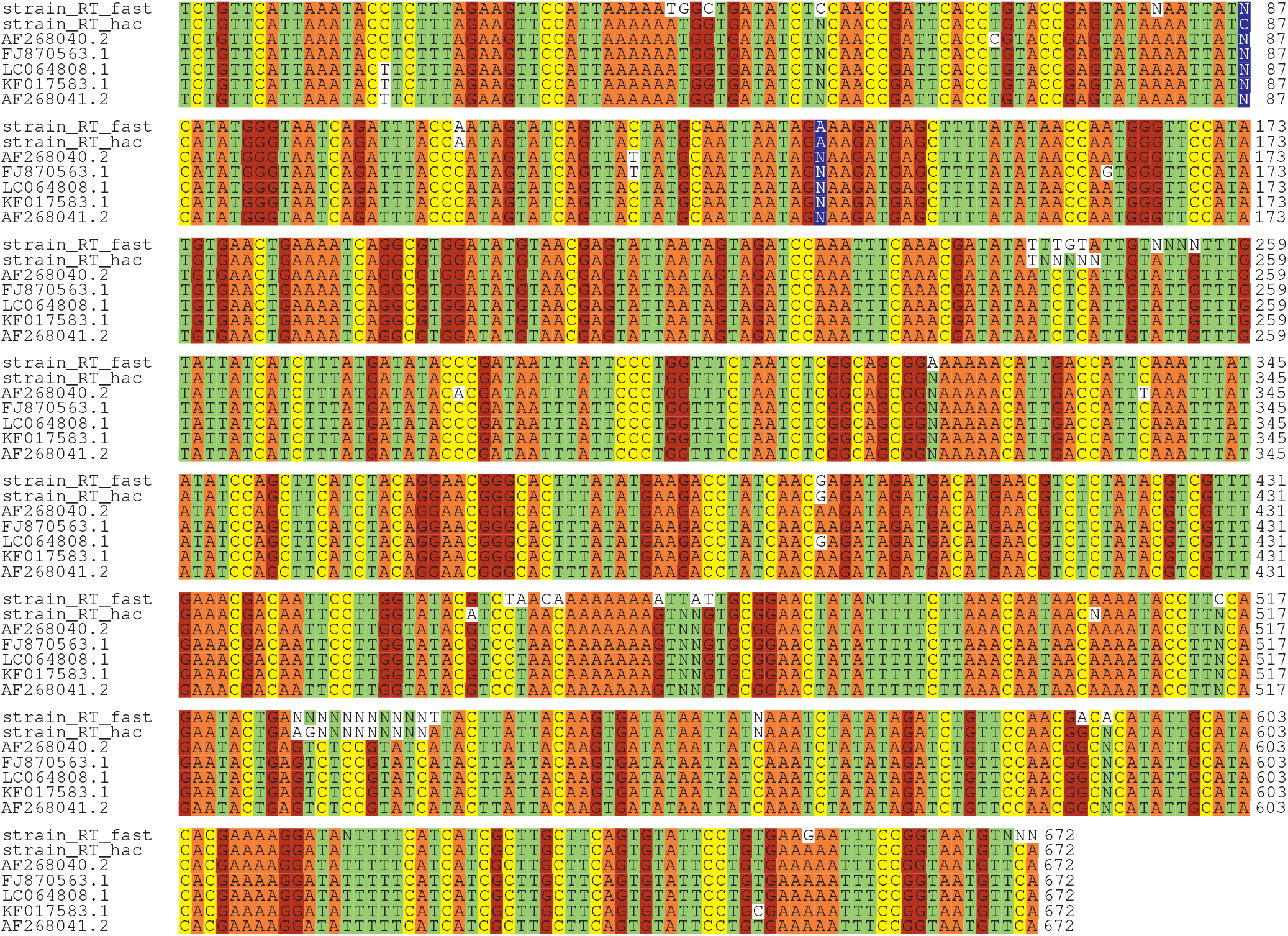
Multiple sequence alignments with the most similar strains. The sequence matching gene *gB* basecalled with fast and high-accuracy configurations is represented by label `strain_RT_fast` and `strain_RT_hac`, respectively. Strain `FJ870563.1` and `KF017583.1` (representative reference genome) strains originated from China, `AF268041.2` and `LC064808.1` from Japan, while the `AF268040.2` one from Spain. The unique insertions are highlighted by blue.

## Discussion

By an ONT-based metagenomic study, we identified sequences of PCMV, including a 650 bp long one that matched the *gB* gene. It is the second report of the virus presence in Hungary but the first with a comparable genomic sequence.

High-accuracy basecalling resulted in fewer polymorphisms and shorter deletions compared to fast basecalling. Although we cannot know with certainty the exact sequence of the virus in our sample, we suppose that polymorphisms identified by the high-accuracy approach may be reliable. Not only do the deletions represent a difference from the reference genome, but none of the *gB* genes has similar ones. A closer look at the pattern of the reference genome at the beginning of the deletions in Figure 3 reveals a sequence `TCTC` . This may be a short tandem repeat (STR or microsatellite), which Delahaye and Nicolas^18^ associate with the appearance of deletions originating from the ONT-basecall. Unfortunately, only one read in our samples matched the *gB* gene. Perhaps if we had sequenced the samples more deeply and had more overlapping reads, these deletions could have been filled in.

The vast majority of PCMV sequences were found in the piglet sample, which the age-specificity of infection and disease can explain. A persistent but mild problem on the farm for several years is the presence of sneezing and rhinitis in piglets up to 6 weeks of age. Several pathogens have been identified in the past to investigate the background of this problem, but their treatment has not resulted in a solution. The small number of reads in the fattening pigs may be a consequence of barcode cross-talk and are, in fact, derived from the piglet sample.^19^ However, it is also possible that minimal levels of the virus are present in the fatteners. This is supported by our metagenomic analysis of nasal swabs from fattening pigs on the same farm, using Illumina sequencing in 2019, in which two pooled samples were analyzed. One had 205, and the other 118 reads matching the PCMV reference genome, but no reads for the *gB* gene.

The experience of our study tells us that ONT-metagenomics can be a promising tool for rapidly detecting pathogens in farm animals. However, instead of using multiplex sequencing as we have done, we should consider using smaller, single-use flow cells to sequence samples individually, avoiding the barcode cross-talks.

## Declarations

### Ethics approval and consent to participate

Not applicable.

### Consent for publication

Not applicable.

### Availability of data and material

The datasets used and/or analysed during the current study available from the corresponding author on reasonable request.

### Competing interests

The authors declare that they have no competing interests.

### Funding

The research was supported by the European Union’s Horizon 2020 research and innovation program under Grant Agreement No. 874735 (VEO).

### Author contributions statement

NS takes responsibility for the data’s integrity and the data analysis’s accuracy. AGT, NS, and TR conceived the concept of the study. AGT, LM, NS, RF, and TR performed sample collection. AGT, ÁB, RF, and SS did the DNA extraction and metagenomics library preparation. NS participated in the bioinformatic analysis. AGT and NS participated in the drafting of the manuscript. AGT, IC, and NS completed the manuscript’s critical revision for important intellectual content. All authors read and approved the final manuscript.

## Notes

### Competing Interest Statement

The authors have declared no competing interest.

